# Stable interaction of NONO with DBHS family members upon etoposide-induced DNA damage

**DOI:** 10.1101/2025.03.09.642258

**Authors:** Barbara Trifault, Stephanie Lamer, Andreas Schlosser, Kaspar Burger

## Abstract

The Non-POU domain containing octamer binding (NONO) protein is a member of the multifunctional *Drosophila* behavior/human splicing (DBHS) protein family and a core component of nuclear paraspeckles. NONO forms dimers with the other two DBHS members splicing factor proline and glutamine rich (SFPQ) protein or the paraspeckle component 1 (PSCP1) to modulate RNA metabolism and gene expression both at the transcriptional and post-transcriptional level. Increasing evidence suggests that NONO participates in genome maintenance by stimulating the DNA damage response (DDR) upon induction of DNA double-strand breaks (DSBs). However, the molecular principles that engage NONO in genome stability are poorly understood. We hypothesized that the induction of DSBs alters NONO protein-protein interactions and applied label-free mass spectrometry to test for changes in the interactome of NONO in human U2OS cells upon treatment with the topoisomerase II inhibitor etoposide. Surprisingly, our mass spectrometry data reveal that etoposide treatment does not induce major changes in NONO protein-protein interactions. We confirmed this finding by orthogonal co-immunoprecipitation assays and co-localization assays. Our data suggest that the bulk of interactions between NONO and SFPQ or PSPC1 are insensitive to etoposide treatment and that DBHS family members promote genome stability as stable dimers.

## INTRODUCTION

RNA-binding proteins (RBPs) are an abundant protein family with critical roles in RNA metabolism. Utilizing a characteristic RNA recognition motif (RRM) and variable RNA-binding domains, RBPs recognize a diverse array of transcripts to participate in virtually all aspects of gene expression, ranging from the production and processing of nascent RNA in the nucleus to posttranscriptional roles in translation and RNA stability [1,2]. The Non-POU domain containing octamer binding (NONO) protein is a member of the *Drosophila* behavior/human splicing (DBHS) protein family and concentrates in paraspeckles. Paraspeckles are stress-responsive nuclear bodies that are built on the lncRNA NEAT1 and sequester 40 or so RBPs to form RNA metabolic hubs [3–7]. Members of the DBHS family are abundant and dynamic RBPs that mediate numerous protein-protein and protein-nucleic acid interactions to participate in various steps of pre-mRNA synthesis [8]. NONO self-dimerizes or forms heterodimers with the splicing factor proline and glutamine rich (SFPQ) protein or the paraspeckle component 1 (PSCP1) protein to promote the retention and editing of a subset of pre-mRNA transcripts within paraspeckles [9,10]. NONO also enriches on chromatin and associates with promoter-associated nascent transcripts to stimulate the formation of transcriptional condensates and augment RNA polymerase II activity [11–15]. First discovered as a splicing factor, NONO directly interacts with spliceosomal subunits and recognizes the 5’ splice site, but is not essential for the splicing machinery in unperturbed cells [16,17]. Instead, NONO enhances pre-mRNA processing of super enhancer-associated transcripts upon oncogenic stress [18]. NONO is upregulated in many tumors and high levels of NONO foster the expression oncogenic transcriptional programs, which promotes proliferation and drug-resistance of cancer cells [19]. Thus, NONO is a multifunctional RNA metabolic factor.

Genome instability is a hallmark of cancer [20]. DNA double-strand breaks (DSBs) are highly toxic and threat genome stability [21,22]. To counteract genotoxic stress, the DNA damage response (DDR) engages a myriad of proteins that sense and repair DSBs, including RBPs [23–25]. NONO and other DBHS family members are also linked to DDR and modulates the recognition and repair of DSBs [26]. Early studies suggest that NONO heterodimerizes with SFPQ to promote DSB pathway choice [27]. The depletion of NONO indeed delays the clearance of DSB foci and increases chromosomal aberrations [28,29]. NONO directly accumulates at DSBs and interacts with a subset of DDR factors, which fosters DSB repair via non-homologous end-joining (NHEJ) [30,31]. The accumulation of NONO at DSBs occurs in a poly(ADP-ribose)-dependent manner and involves the activation of the DSB-sensing DNA-dependent protein kinase (DNA-PK) [28,32]. More recently, novel mechanisms for the genome protective role of NONO have been reported, which include the relocalization of NONO and other DBHS family members to the nucleolus and nucleolus-associated condensates via interaction with nucleolar transcripts [33,34]. These findings underscore the relevance of NONO in promoting efficient DSB repair and genome stability.

Interestingly, NONO binding affinity to RNA substrates is regulated by the coactivator associated arginine methyltransferase 1 (CARM1/PRMT4) [35]. Moreover, NONO undergoes glycosylation and ubiquitinylation to efficiently accumulate at DSBs and UV-induced lesions, respectively [36,37]. However, to what extent DNA damage impacts on NONO protein-protein interactions is poorly understood. We hypothesized that the induction of DSBs alters the interactions of NONO with other DBHS family proteins and employed label-free quantitation by mass spectrometry to determine NONO protein-protein interactions in human U2OS cells in the absence or presence of the topoisomerase II inhibitor etoposide. Surprisingly, we observed no clear changes in the NONO interactome and persistent association of both SFPQ and PSPC1 to NONO. Persistent interaction of NONO with DBHS family members in response to etoposide treatment was validated by orthogonal assays and suggests remarkable stability of the bulk of *bona fide* NONO protein-protein interactions upon induction of DSB signaling.

## MATERIALS AND METHODS

### Tissue culture, transfections, inhibitors and antibodies

Human U2OS osteosarcoma cells were acquired from the German collection of microorganisms and cell cultures (DSMZ), cultured at 37°C and 5 % CO2 in Dulbecco’s modified eagle’s medium (DMEM, Gibco) complemented with 10 % fetal bovine serum (FBS, Capricorn), 100 U/mL penicillin-streptomycin (Gibco) and 2 mM L-glutamine (Gibco), and routinely tested for mycoplasma infection. For ectopic expression, cells were transfected with 1.5 µg plasmid DNA encoding HA-tagged NONO (pcDNA3.1-HA-NONO, kind gift from Nicolas Manel). For knockdown experiments, cells were transfected with 100 nM siRNA that selectively targets either NONO (siNONO, Dharmacon, L-007756-01 smart pool) or represents non-targeting control (siControl, Dharmacon, D-001810-01 smart pool). Lipofectamine 2000 (Invitrogen) and Opti-MEM (Gibco) was used for transfections according to the manufactureŕs protocol. Incubation conditions with etoposide (Sigma) were for 2 h with 20 µM, unless stated differently. The following antibodies were used in this study. Primary antibodies were: IgG control (Proteintech, 30000-0-AP); NONO (Proteintech, 11058-1-AP); PSPC1 (Proteintech, 16714-1-AP); SFPQ (Abcam, ab177149); Ki-67 (Thermo, MA5-14520); HA-tag (Abcam, ab9110; BioLegend, 901502); Fibrillarin (Abcam, ab5821); Vinculin (Sigma, V9131); histone H2B (Abcam, ab1790); phospho-ser-139 histone H2A.X (Cell Signaling, 2577; Millipore, 05- 636); 53BP1 (Novus, NB100-304). Secondary antibodies were: HRP-linked IgG (Cytiva, GEHENA934; Cytiva, NA931); Alexa Fluor 546-linked IgG (Thermo, A10036), Alexa Fluor 488-linked IgG (Thermo, A21206), Alexa Fluor 546-linked IgG (Thermo, A10040).

### Immunoblotting and immunoprecipitation

Cells were lysed, boiled (5 min, 95°C) and sonicated (Branson, 10 %, 5 sec) in 4x sample buffer (250 mM tris-HCl pH 6.8, 8 % sodium dodecyl sulphate (SDS), 40 % glycerol, 0.8 % β- mercaptoethanol, 0.02 % bromophenol blue). Whole cell extracts were separated by denaturating SDS polyacrylamide gel electrophoresis (PAGE) and transferred onto a nitrocellulose membrane (Cytiva). Membranes were blocked (30 min, RT) in PBST (PBS/0.1 % triton x-100) with 5 % milk, probed with primary (overnight, 4°C) and secondary antibodies (1 h, RT) diluted in PBST with or without milk and washed in between incubation steps with PBST. Signals were visualized with an ECL kit (Cytiva) with an imaging station (Fusion FX Vilber, LAS-4000 Fuji) and quantified with ImageJ (NIH). For immunoprecipitation, cells were trypsinised, resuspended in DMEM, centrifuged (1200 rpm, 3 min), washed in cold PBS and centrifuged again. The pellet was lysed for 10 min on ice in 5 volumes of IP buffer (200 mM NaCl, 0.5 mM EDTA, 20 mM HEPES, 0.2 % NP-40, 10 % glycerol, 1x protease/phosphatase inhibitor (Roche), 100 U of Ribolock RNase inhibitor (Thermo). Lysates were centrifuged (12000 rpm, 12 min) and supernatants were collected in a new tube. 10 % of lysate was kept as input (IN) and the remaining 90 % was incubated (3 h, 4°C) on a rotative wheel with 5 µg primary antibody pre-conjugated with 30 µL G beads overnight. Crosslinking of pre-conjugated antibodies was performed with bis(sulfosuccinimidyl)suberate (BS3, Thermo) or dimethyl pimelimidate (DMP, Thermo) according to the manufacturer’s protocol. Immunocomplexes were captured on a magnet (Invitrogen), washed three times with 800 µL IP buffer and eluted (5 min, 95°C) with 40 µL of 4x sample buffer (reducing condition). For native PAGE, samples were eluted in 4x sample buffer without β-mercaptoethanol (non-reducing condition). Proteins were visualized by using either a SilverQuest staining kit (Thermo) according to the manufactureŕs protocol prior to transfer or by staining for 5 min at RT with Ponceau solution (0.5 % Ponceau S/1 % acetic acid) after the transfer.

### Confocal imaging

Cells grown on cover slips (Roth) were washed in PBS, fixed (10 min, RT) with 3 % paraformaldehyde (Sigma), permeabilized (10 min, RT) with PBST, washed (5 min, three times) in PBS and blocked (2 h, 4°C) with PBS/10 % FBS. For staining, cells were incubated in a humidified chamber with primary antibodies (overnight at 4°C) and secondary antibodies (2 h, RT) diluted in PBS/0.15 % FBS and secondary antibodies. Cells were washed with PBST (three times, 5 min) between incubations and sealed in 4,6-diamidino-2-phenylindole (DAPI)- containing mounting medium (Vectashield). Signals were imaged with a confocal microscope (CLSM-Leica-SP2, x63 objective, 1024×1024 resolution, airy = 1) and channels were acquired sequentially, between frames, with equal exposure times as single scans. Images were analyzed and co-localization was assessed by ImageJ and RGB profiler (NIH).

### Nanoscale liquid chromatography coupled to tandem mass spectrometry (Nano LC-MS/MS)

For in-gel digestion, immunoselected proteins were eluted with 4x lithium dodecyl sulphate (LDS) sample buffer (NuPAGE, Thermo) according to the manufacturer’s protocol and precipitated in a four-fold volume of acetone (overnight, −20°C). Precipitated proteins were washed three times with acetone at −20°C, dissolved in 4x LDS sample buffer, reduced (10 min, 70°C) with 50 mM DTT and alkylated (20 min, RT) with 120 mM iodoacetamide. Proteins were separated by Novex 4-12 % bis-tris gels with MOPS buffer (NuPAGE, Thermo) according to the manufacturer’s protocol. Gels were washed (5 min, three times) with ultrapure water, stained (60 min, RT) with Coomassie (Thermo) and washed again for 1 h. All gel lanes were cut in 15 slices, destained with 30 % acetonitrile in 0.1 M NH_4_HCO_3_ pH 8.0, shrunk with 100 % acetonitrile, and dried in a vacuum concentrator. Each gel band was digested (overnight, 37°C) with 0.1 µg trypsin in 0.1 M NH_4_HCO_3_ pH 8.0, the supernatant was removed and the peptides were extracted from the gel slices with 5 % formic acid. The extracted peptides were pooled with the supernatant. For NanoLC-MS/MS analysis, an Orbitrap Fusion (Thermo) equipped with a PicoView Ion Source and coupled to an EASY-nLC 1000 was used. Peptides were loaded on capillary columns (PicoFrit, 30 cm x 150 µm ID, New Objective) self-packed with ReproSil-Pur 120 C18-AQ, 1.9 µm (Dr. Maisch) and separated with a linear gradient from 3 % to 30 % acetonitrile and 0.1 % formic acid (30 min, flow rate 500 nL/min). Scans were acquired with a resolution of 7500 for MS/MS scans and 60000 for MS scans in the Orbitrap analyzer. HCD fragmentation with 35 % normalized collision energy was applied and a top speed data-dependent MS/MS method with a fixed cycle time of 3 sec was used. Dynamic exclusion was set with a repeat count of 1 and an 30sec exclusion duration and singly charged precursors were excluded from selection. Minimum signal threshold for precursor selection was set to 5×104 and predictive AGC was used with a AGC target value of 2×105 for MS scans and 5×104 for MS/MS scans. EASY-IC was used for internal calibration.

### Mass spectrometry data analysis

Raw MS data files were analyzed with MaxQuant version 1.6.2.2 [38]. Database search was performed with Andromeda, which is integrated in the utilized version of MaxQuant. The search was performed against the UniProt Human Reference Proteome database (January 2021, UP000005640, 75777 entries). Additionally, a database containing common contaminants was used. The search was performed with tryptic cleavage specificity with 3 allowed miscleavages. Protein identification was monitored by setting a false-discovery rate (FDR) of <1 % on protein and peptide spectrum match level. In addition to MaxQuant default settings, the search was performed against the following variable modifications: Protein N-terminal acetylation, Gln to pyro-Glu formation (N-term. Gln) and oxidation (Met). Carbamidomethyl (Cys) was set as fixed modification. Further data analysis was performed using R scripts developed in-house. LFQ intensities were used for protein quantitation [39]. Proteins with less than two razor/unique peptides were removed. Missing LFQ intensities were imputed with values close to the baseline. Data imputation was performed with values from a standard normal distribution with a mean of the 5 % quantile of the combined log10-transformed LFQ intensities and a standard deviation of 0.1. For the identification of significantly enriched proteins, median log2 transformed protein ratios were calculated from the two replicate experiments and boxplot outliers were identified in intensity bins of at least 300 proteins. Log2 transformed protein ratios of sample versus control with values outside a 1.5x (significance 1) or 3x (significance 2) interquartile range (IQR), respectively, were considered as significantly enriched in the individual replicates.

### Ethics statement

All data are derived from the U2OS cancer cell line (non-pathogenic, non-infectious, safety level S1), which has been isolated in 1964. U2OS cells are an established cancer model cell line for more than 50 years and working with U2OS cells does not require ethical approval.

## RESULTS

### Validation of NONO antibody from mass spectrometry

We wished to study the impact of DNA damage on the NONO interactome in U2OS cells. We acquired a knockout-validated polyclonal NONO antibody and confirmed its selectivity upon RNAi-mediated depletion of NONO. Using the NONO antibody for immunoblotting and immunofluorescence microscopy, we observed prominent reduction in reactivity upon transfection of NONO-selective siRNA (Fig. S1A-C). Next, we validated the induction of DNA damage by confocal imaging of DSB markers in response to etoposide treatment. As expected, we observed prominent formation of nuclear foci in the presence of etoposide that stained positive for the *bona fide* DSB marker p53-binding protein 1 (53BP1) and partially overlapped with signals for the DNA damage marker serine-139 phosphorylated histone H2A.X variant (γH2A.X) (Fig. S1D and E). We also tested the capacity of the antibody to immunoselect NONO and performed immunoprecipitation experiments (Fig. S1F). Indeed, the antibody efficiently enriched NONO from U2OS whole cell lysates, irrespective of etoposide treatment conditions. Thus, we established conditions to detect and immunoselect NONO in the absence and presence of DSBs.

We noticed prominent immunoglobulins signals in our immunoprecipitation samples, suggesting prominent coelution of bead-conjugated immunoglobulins. Excessive amounts of immunoglobulins can perturb mass spectrometry data and prevent the detection of less abundant peptides. Thus, aimed to covalently bind the antibody to the matrix and prevent coelution. We elaborated on our immunoprecipitation protocol and introduced a crosslinking step using either BS3 or DMP reagent (Fig. S1G). Using non-reducing conditions, we found that both BS3 and DMP efficiently prevented coelution of immunoglobulins from the beads. However, the NONO antibody only tolerated BS3, but not DMP crosslinking and the latter prevented the efficient immunoselection of NONO from lysates. We repeated the immunoprecipitation experiment and assessed NONO levels after either non-reducing or reducing separation conditions (Fig. S1H). Immunoblotting for NONO confirmed that BS3 crosslinking both preserves the affinity of the antibody toward NONO and prevents coelution of immunoglobulins from the beads. We conclude that etoposide treatment and antibody crosslinking by BS3 are appropriate to assess changes in the NONO interactome upon DNA damage.

### Persistent interaction of NONO with DBHS family members upon DNA damage

Having established conditions to immunoselect NONO, we applied mass spectrometry to determine the impact of DNA damage on NONO protein-protein interactions. For comparative analysis of two biological replicates, we incubated cells in the absence or presence of DNA damage and performed immunoprecipitation from whole cell lysates using either the NONO antibody or an isotype control (Fig. S2A). Correlation analysis revealed a high similarity between replicates, irrespective of the antibody used for immunoprecipitation or etoposide treatment (Fig. S2B). Overall, we detected 1031 proteins that were reproducibly present upon NONO immunoprecipitation, albeit many of them with low peptide counts or poor sequence coverage. To increase stringency, we considered only those as hits that were detected by at least five unique peptides in the NONO immunoprecipitation samples. Such filtering narrowed down the list of proteins to 238 that were immunoselected by the NONO antibody in the absence or presence of etoposide treatment (Fig. 1A). Reassuringly, NONO was found as top hit together with SFPQ and PSPC1, and all three DBHS family members scored with high significance (Table S1). However, visual inspection of mass spectrometry data suggested an overall high similarity between control and DNA damage conditions. Indeed, a stable, etoposide-resistant interaction between NONO and DBHS family members was confirmed by immunoblotting (Fig. 1B and C). Also, 76 of the top100 mass spectrometry hits were found in both conditions (Fig. 1D; Table S2). To directly assess for differential co-enrichment of proteins in the absence or presence of DNA damage, we integrated both treatment conditions into one graph (Fig. 2A). Linear regression analysis revealed that PSPC1, SFPQ and virtually all other hits were co-immunoprecipitated with NONO irrespective of etoposide treatment. We considered a second filter and scored only those factors as significant that were at least 4-fold enriched over the isotype control. This analysis yielded 49 proteins, out of which 28 were reported as nuclear proteins. With a few exceptions, the list of top30 proteins is strikingly similar and the bulk of filtered hits in the control condition are also found upon etoposide treatment (Fig. 2B). Of note, 90% or so of hits scoring as nuclear proteins were RBPs (Fig. S2C) and these included several previously reported components of paraspeckles and NEAT1 interactors, such as HNRNPUL1, YBX1 or SYNCRIP [40,41]. We conclude that our NONO interactome data are valid, but may suffer from a considerable number of false-positive hits and cytoplasmic contaminants. Surprisingly, the comparative analysis of NONO interactomes revealed no significant changes in response to DNA damage.

**Figure 1.**
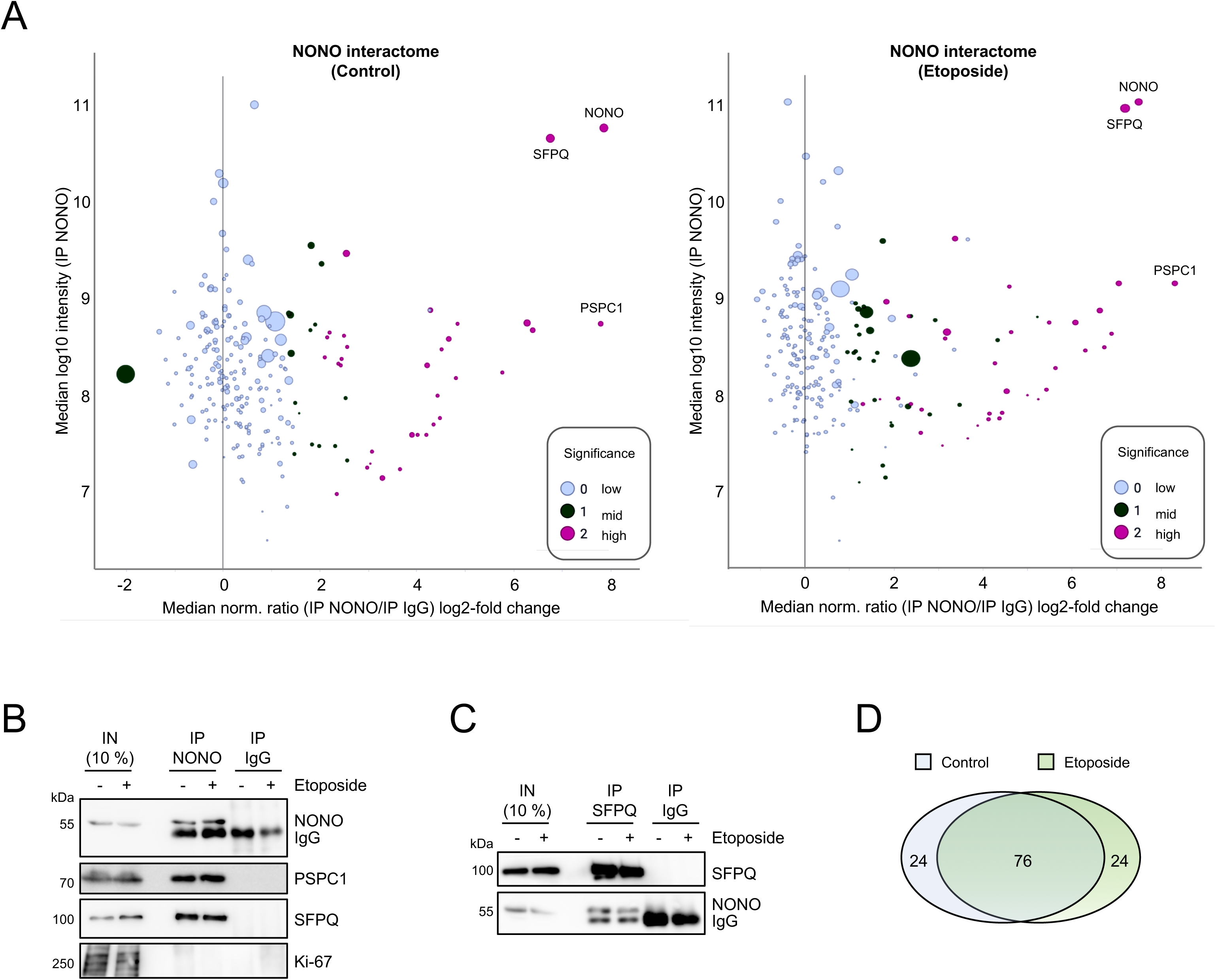
Assessment of NONO interactomes in U2OS cells. (A) Intensity versus ratio plots displaying NONO mass spectrometry hits. 238 proteins that were detected with at least five unique peptides identified by mass spectrometry are shown. Data are normalized to IgG isotype controls and represent two merged biological replicates. The x-axis/y-axis displays ratios/intensities of peptides. The number of unique peptides correlates with the size of each circle and the significance is color-coded. (B, C) Immunoblots detecting NONO, PSPC1 and SFPQ in whole cell extracts (IN, input) and upon immunoprecipitation (IP) with NONO antibody (B) or SFPQ antibody (C). IgG, isotype control; Ki-67, negative control. (D) Venn diagram showing overlap of NONO mass spectrometry hits. The top100 proteins have been assessed.

**Figure 2.**
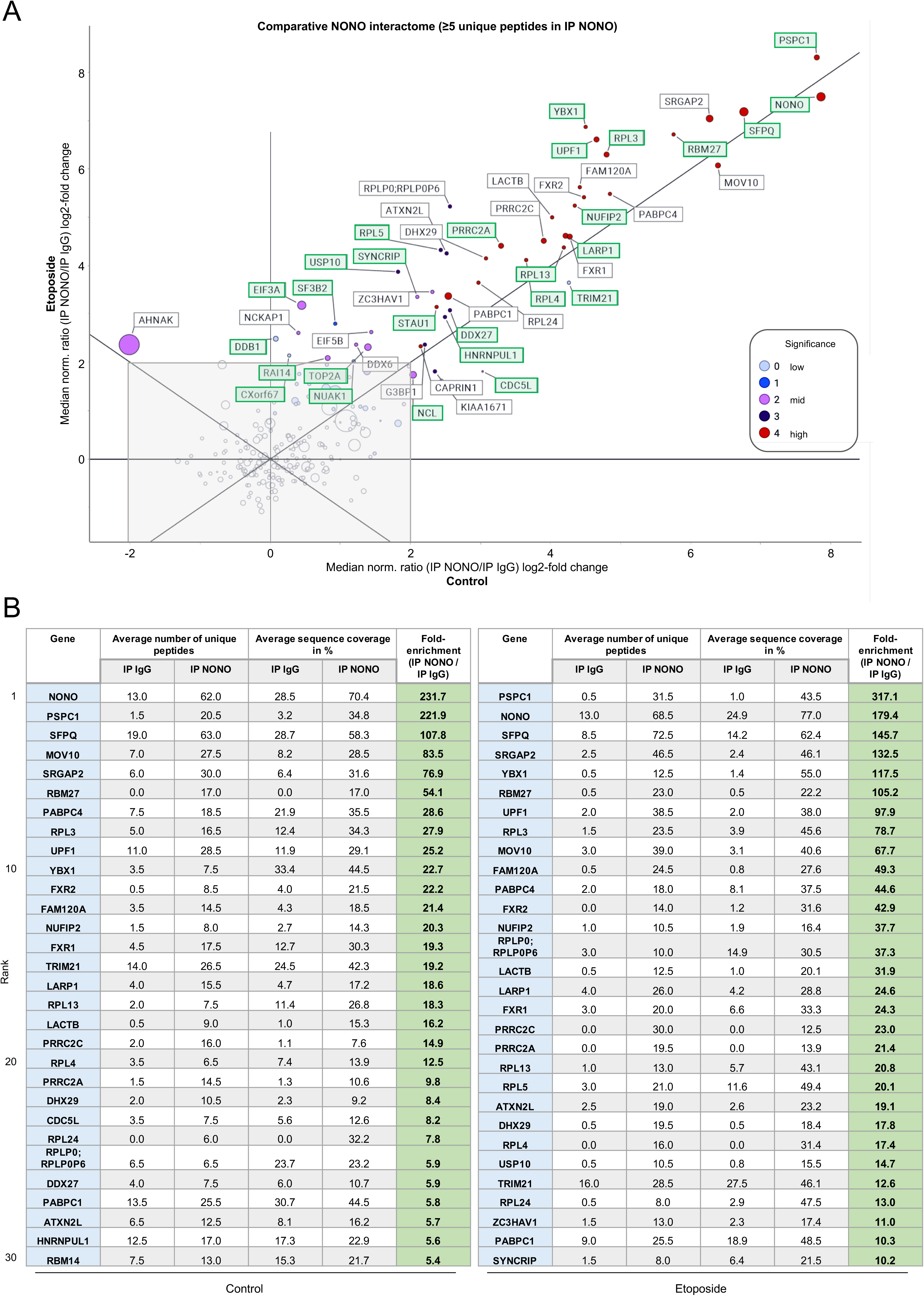
Comparative analysis of NONO mass spectrometry hits in U2OS cells. (A) Integration of mass spectrometry conditions by ratio versus ratio depiction. 49 proteins that were detected with at least five unique peptides identified by mass spectrometry and that were at least 4-fold enriched over IgG isotype controls are shown. Data represent two merged biological replicates. Both the x-axis and the y-axis display ratios of peptides, whilst the diagonal axis displays intensities. The number of unique peptides correlates with the size of each circle and the significance is color-coded. Green box, nuclear protein; grey box, non-significant hits. (B) Table summarizing mass spectrometry data for top30 proteins in the absence or presence of etoposide.

### Co-localization of DBHS family members *in vivo* and concomitant nucleolar re-localization upon DNA damage

We could previously show that NONO, SFPQ and PSPC1 undergo nucleolar re-localization upon etoposide treatment [34,42]. To corroborate our biochemical data and further assess the association of NONO with DBHS family members in response to DNA damage *in vivo*, we investigated the subcellular localization of DBHS proteins in the absence and presence of etoposide by confocal imaging. We aimed to co-stain endogenous SFPQ or PSPC1 with endogenous NONO using the same NONO antibody as used for biochemical assays. To the best of our knowledge, however, there are currently no suitable and validated SFPQ or PSPC1 antibodies available that would allow co-staining with this NONO antibody. Thus, we ectopically expressed full length, HA-tagged NONO (HA-NONO), in U2OS. As expected, HA- NONO localized to the nucleoplasm in unperturbed cells and displayed a more pan-nuclear localization upon etoposide treatment (Fig. S3A). A fraction of HA-NONO also accumulated in the nucleolus, as monitored by partial co-localization with the nucleolar marker fibrillarin (Fig. S3B). Importantly, treatment with etoposide did not alter the expression levels of HA- NONO (Fig. S3C). Next, we co-stained endogenous DBHS proteins with HA-NONO. Reassuringly, HA antibody reactivity displayed strong co-localization with each of the three DBHS family members irrespective of etoposide treatment (Fig. 3A and B). Moreover, pan-nuclear localization could be observed for a fraction of both HA-NONO, and each of the three endogenous DBHS proteins in response to etoposide treatment. We conclude that NONO associates with SFPQ and PSCP1 *in vivo* and that the bulk of these associations is resistant to etoposide treatment.

**Figure 3.**
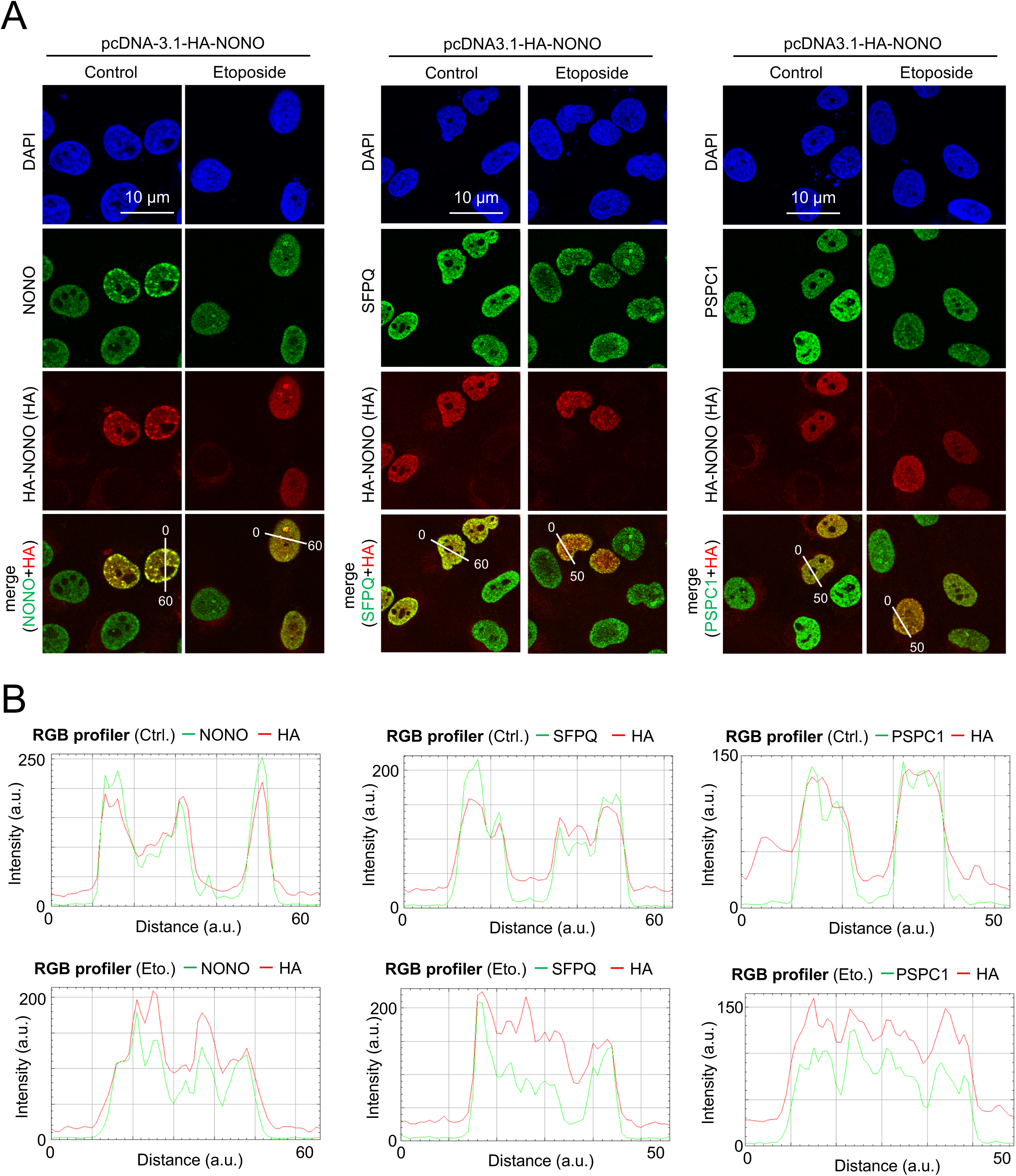
Assessment of DBHS protein localization in U2OS cells. (A) Confocal imaging of either NONO (left), SFPQ (middle) or PSPC1 (right) in combination with HA-NONO upon transient transfection of pcDNA3.1-HA-NONO expression plasmid. Representative images are shown. (B) Quantitation of (A) using RGB profiler line scan analysis. a.u., arbitrary units.

## DISCUSSION

Our study compared NONO protein interactions in the absence or presence of DNA damage. Our mass spectrometry data confirmed the interaction of NONO with DBHS family member, which was further validated by co-immunoprecipitation assays and co-localization studies. The interactome data further indicate that NONO associates with numerous other RBPs. However, our approach did not reveal any clear change in NONO interactions upon induction of DSBs. Albeit confirming stable, etoposide-resistant formation of dimers between NONO and other DBHS family members, the interpretation of DNA damage-induced changes in NONO interactions may suffer from limitations.

NONO shares more than 70 % of sequence homology with SFPQ and PSPC1. The conserved DBHS region of NONO contains two RNA recognition motifs (RRM1 and RRM2), a NONA/paraspeckle domain (NOPS), and a coiled-coil domain [8]. NONO also contains an amino-terminal histidine proline glutamine (HPQ)-rich region and a carboxy-terminal glutamine proline (GP)-rich region, which harbors a nuclear localization sequence (NLS). The canonical RRM1 and non-canonical RRM2 facilitate the binding of NONO to various nucleic acid substrates, whereas the dimerization of DBHS proteins relies on the interaction between the RRM2 domain of one paralog with the NOPS and coiled-coil domains of the other paralog [8,43]. Intriguingly, phosphoproteomic data suggest that the induction of DNA damage by ionizing radiation alters phosphorylation at NONO at threonine residues T428 and T450, which are located in the carboxy-terminal region of NONO [44]. It is tempting to speculate that NONO also undergoes differential phosphorylation upon etoposide treatment and that DNA damage-responsive post-translational modifications alter NONO protein-protein interactions. However, NONO is a multifunctional RBPs and modulates numerous processes in the nucleus. Thus, the fraction of NONO susceptible to DNA damage may be rather small and its analysis likely requires the generation of more selective tools such as phospho-specific antibodies. Our imaging data using a pan-NONO antibody indeed indicate that a minor fraction of the nuclear NONO pool is responsive to etoposide treatment, at least for the conditions used in this study. Typically, factors involved in the recognition and repair of DSBs accumulate at such lesions and form foci that can be visualized by microscopy. However, we did not observe the formation of NONO-positive foci. A selective phospho-specific NONO antibody may not only reveal the formation of NONO-positive DSB foci, but also facilitate a refined analysis of the NONO protein-protein interactome in response to DNA damage. The data set could further benefit from improvements in the mass spectrometry protocol, such as immunoprecipitation from subcellular fractions and the usage of orthogonal assays to study interactomes, such as proximity labelling approaches. The latter allows profiling of proteins that localize in distinct subcellular compartments including the nucleolus [45]. This may define changes in NONO protein-protein interactions in the future.

Our data indicate that a subset of NONO accumulates in the nucleolus upon etoposide treatment. However, NONO relocalization is not quantitative, and therefore potential changes in NONO interactions upon DNA damage are difficult to resolve by mass spectrometry. Interestingly, a more quantitative relocalization of NONO and other DBHS to the nucleolus has been observed upon ionizing radiation [46]. Thus, DNA damage-induced changes in the NONO interactome may occur concomitant with relocalization, for instance upon harsher treatment conditions that induce more persistent DSBs and likely trigger a more comprehensive engagement of NONO in DDR.

Overall, our study provides insights in the interactome of NONO in response to etoposide treatment and suggests remarkable stability of the bulk of NONO protein-protein interactions upon induction of DNA damage. It will be important to determine if the genome-protective role of NONO occurs mostly in association with other DBHS family members or if distinct types of DNA damage selectively engage individual DBHS protein family members in DDR.

## DATA AVAILABILITY

Proteomics data are available at the ProteomeXchange consortium via the PRIDE partner repository [47] with the identifier PXD061274 (reviewer token: aFmB1YlDmtL1) or upon login to the PRIDE website via reviewer account (username, password: 3Uo7pqlSPyJP).

## CONFLICT OF INTEREST

The authors declare no conflict of interest.

## SUPPORTING INFORMATION

**Figure S1.** Validation of NONO antibody selectivity in U2OS cells. (A, B) Immunoblot detecting NONO in whole cell extracts (A) and confocal imaging of NONO (B) upon transient transfection of siRNA targeting NONO (siNONO) or non-targeting control siRNA (siControl). Vinculin, loading control; broken circle, nucleus. Representative images are shown. (C) Quantitation of (B) using RGB profiler line scan analysis. a.u., arbitrary units. (D) Confocal imaging of 53BP1 and γH2A.X foci. Representative images are shown. (E) Quantitation of (D) using RGB profiler line scan analysis. (F) Immunoblot detecting NONO upon immunoprecipitation (IP) with NONO antibody from whole cell lysates (IN, input). IgG, isotype control antibody. (G) Immunoblot detecting NONO upon immunoprecipitation (IP) with NONO antibody from whole cell lysates and native PAGE. Silver stain, loading control; exp., exposure. Full blots are shown. (H) as in (G) but comparing NONO signals upon native PAGE (left) and reducing PAGE (right). Full blots are shown. Control, non-crosslinked condition, BS3, bis(sulfosuccinimidyl)suberate; DMP, dimethyl pimelimidate.

**Figure S2.** Quality control for NONO mass spectrometry in U2OS cells. (A) Coomassie stain upon immunoprecipitation (IP) with NONO antibody from whole cell lysates. IgG, isotype control antibody. (B) Correlation analysis of mass spectrometry replicates. Both the x-axis and the y-axis display intensities of peptides. r, correlation coefficient. (C) Stratification of top30 mass spectrometry hits.

**Figure S3.** Quality control for ectopic expression of HA-tagged NONO in U2OS cells. (A) Confocal imaging of HA-NONO upon transient transfection of pcDNA3.1-HA-NONO expression plasmid. FBL, Fibrillarin, nucleolar marker. Representative images are shown. (B) Quantitation of (A) using RGB profiler line scan analysis. a.u., arbitrary units. (C) Immunoblot detecting HA-NONO upon transient transfection of pcDNA3.1-HA-NONO expression plasmid in whole cell extracts. Vinculin, loading control; mock, non-transfected control.

**Figure S4.** Raw data. Uncropped immunoblots are shown. Red box, cropped area depicted in figure panels.

**Table S1.** Mass spectrometry data.

**Table S2.** List of top100 mass spectrometry hits.

## ACKNOWLEDGMENTS

We thank all members of the department for support. We acknowledge Lea Boten and Teresa Klugt for excellent technical assistance. This work was funded by a grant from the German Cancer Aid (grant code: 8606100-NG1) awarded to KB.

## Author contributions

Conceptualization, BT, AS and KB; Methodology, BT, SL, AS and KB; Investigation, BT and KB; Formal Analysis, BT, SL, AS and KB; Writing – Original Draft, BT and KB; Writing – Review & Editing, AS and KB; Funding Acquisition, KB; Supervision, AS and KB.

## Notes

### Competing Interest Statement

The authors have declared no competing interest.

### Summary of Updates

This version of the manuscript has been revised to update Supplemental files.

