## Supplemental material for "Stable interaction of NONO with DBHS family members upon etoposide-induced DNA damage"

### Figure S1

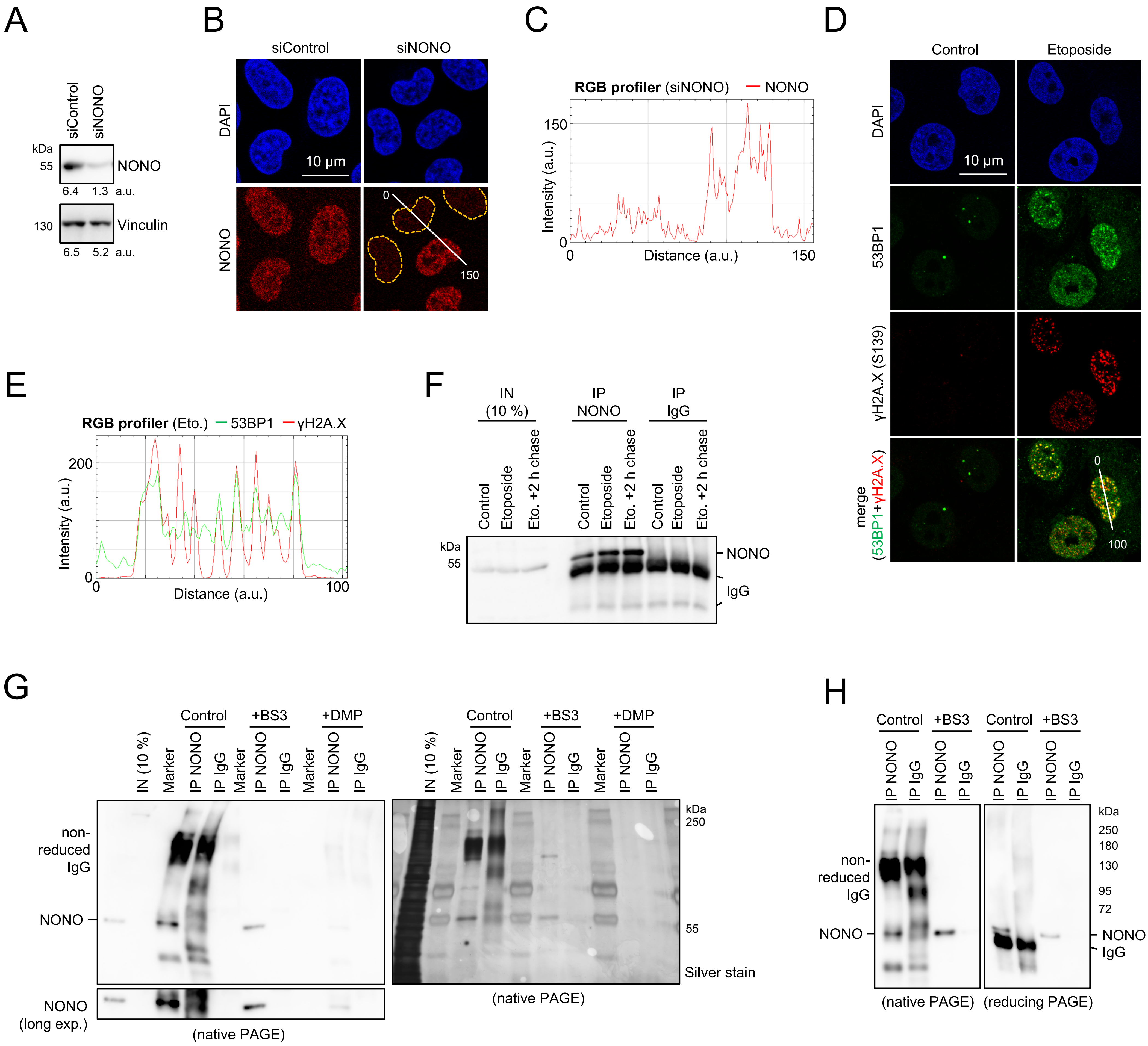


**Figure S1. Validation of NONO antibody selectivity in U2OS cells.** (A, B) Immunoblot detecting NONO in whole cell extracts (A) and confocal imaging of NONO (B) upon transient transfection of siRNA targeting NONO (siNONO) or non-targeting control siRNA (siControl). Vinculin, loading control; broken circle, nucleus. Representative images are shown. (C) Quantitation of (B) using RGB profiler line scan analysis. a.u., arbitrary units. (D) Confocal imaging of 53BP1 and γH2A.X foci. Representative images are shown. (E) Quantitation of (D) using RGB profiler line scan analysis. (F) Immunoblot detecting NONO upon immunoprecipitation (IP) with NONO antibody from whole cell lysates (IN, input). IgG, isotype control antibody. (G) Immunoblot detecting NONO upon immunoprecipitation (IP) with NONO antibody from whole cell lysates and native PAGE. Silver stain, loading control; exp., exposure. Full blots are shown. (H) as in (G) but comparing NONO signals upon native PAGE (left) and reducing PAGE (right). Full blots are shown. Control, non-crosslinked condition, BS3, bis(sulfosuccinimidyl)suberate; DMP, dimethyl pimelimidate.

### Figure S2

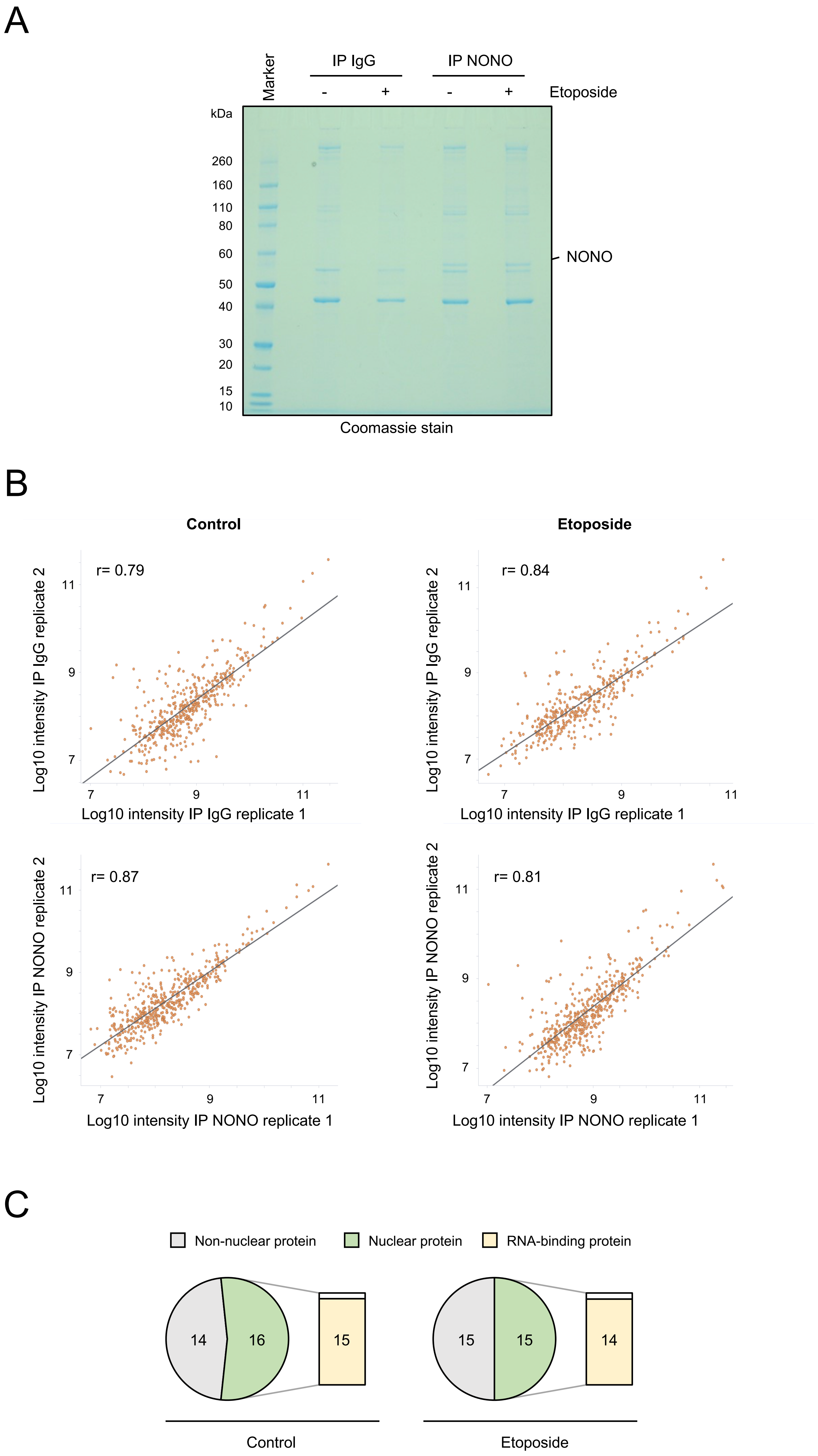


**Figure S2. Quality control for NONO mass spectrometry in U2OS cells.** (A) Coomassie stain upon immunoprecipitation (IP) with NONO antibody from whole cell lysates. IgG, isotype control antibody. (B) Correlation analysis of mass spectrometry replicates. Both the x-axis and the y-axis display intensities of peptides. r, correlation coefficient. (C) Stratification of top30 mass spectrometry hits.

### Figure S3

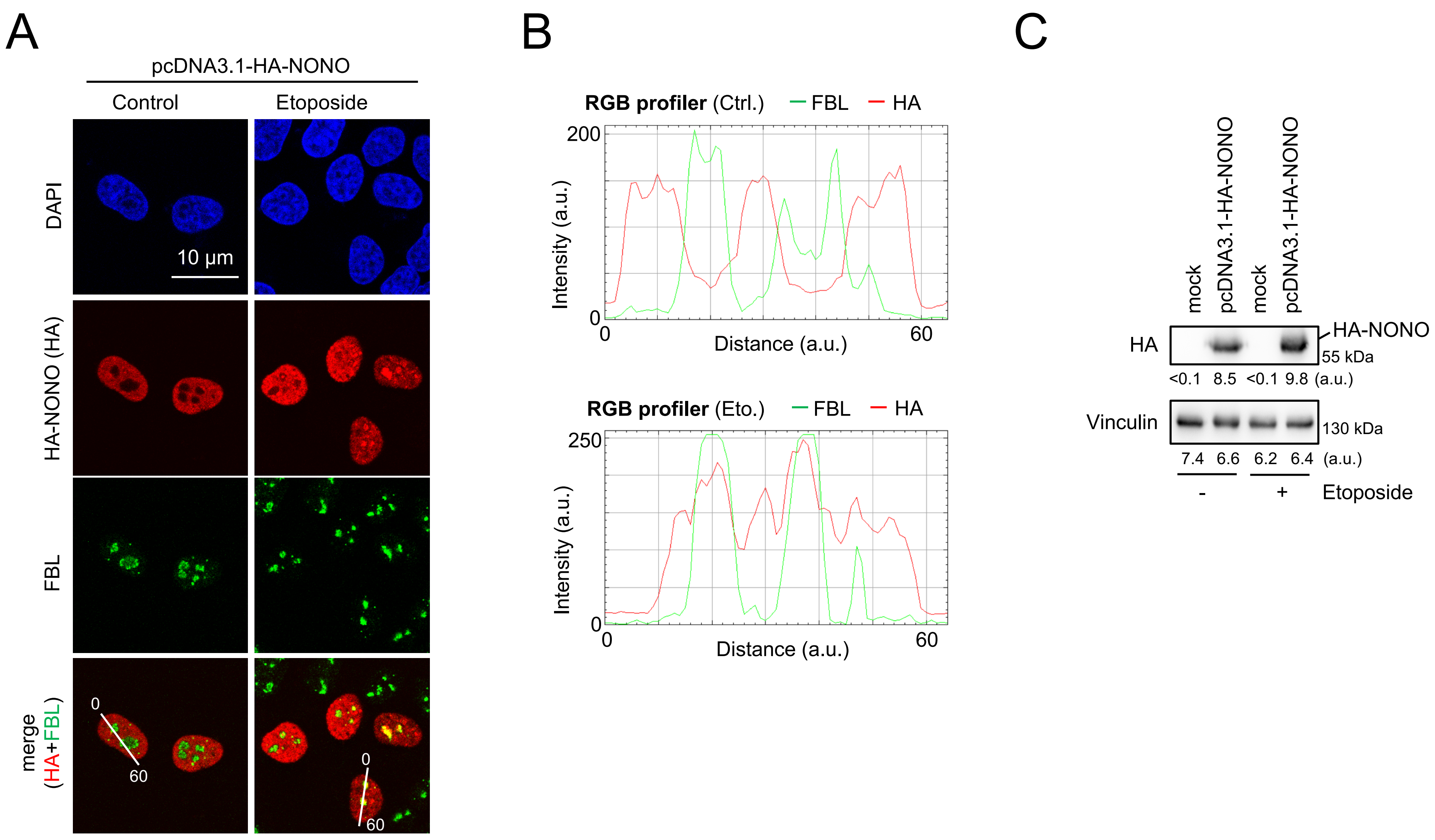


**Figure S3. Quality control for ectopic expression of HA-tagged NONO in U2OS cells.** (A) Confocal imaging of HA-NONO upon transient transfection of pcDNA3.1-HA-NONO expression plasmid. FBL, Fibrillarin, nucleolar marker. Representative images are shown. (B) Quantitation of (A) using RGB profiler line scan analysis. a.u., arbitrary units. (C) Immunoblot detecting HA-NONO upon transient transfection of pcDNA3.1-HA-NONO expression plasmid in whole cell extracts. Vinculin, loading control; mock, non-transfected control.

### Figure S4

#### related to Fig. 1B

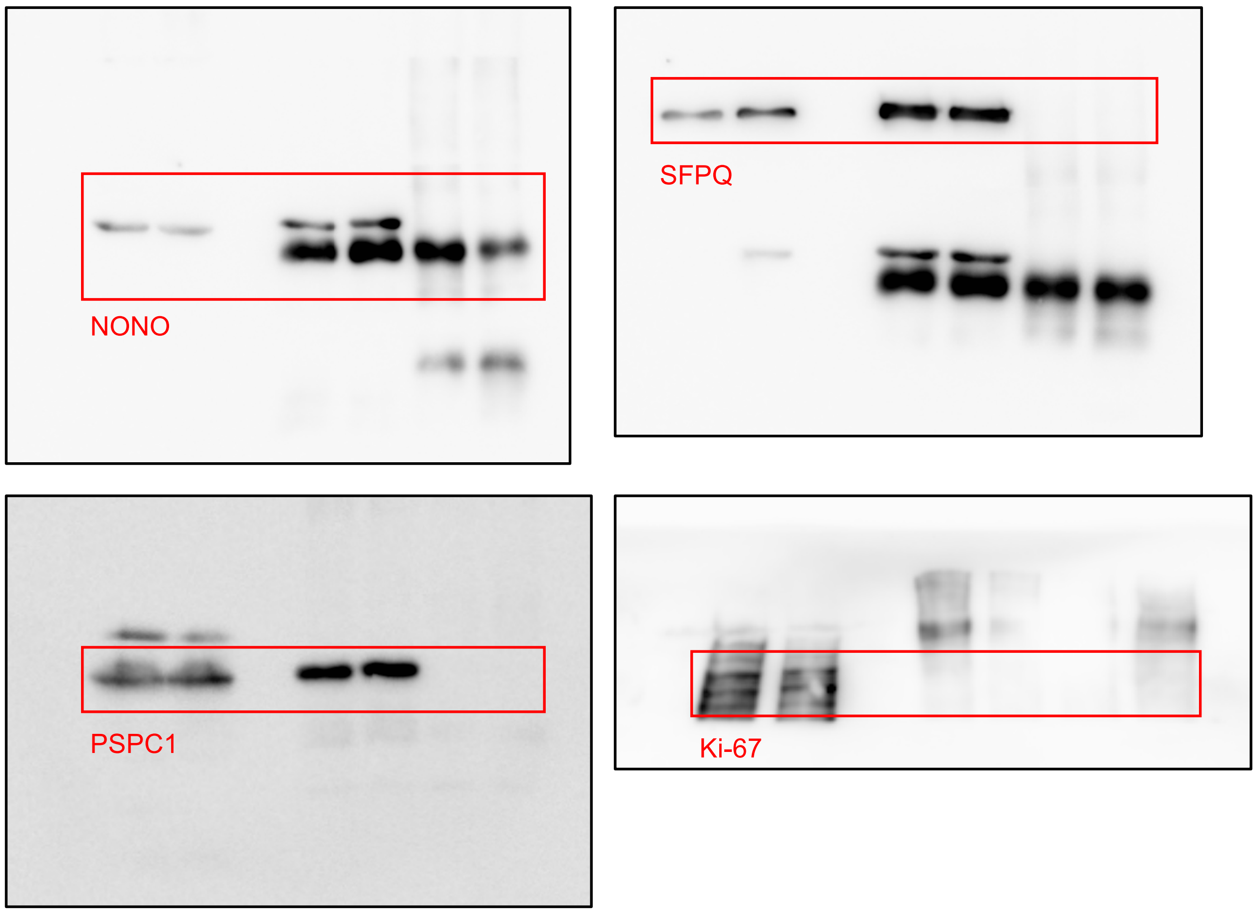


#### related to Fig. 1C

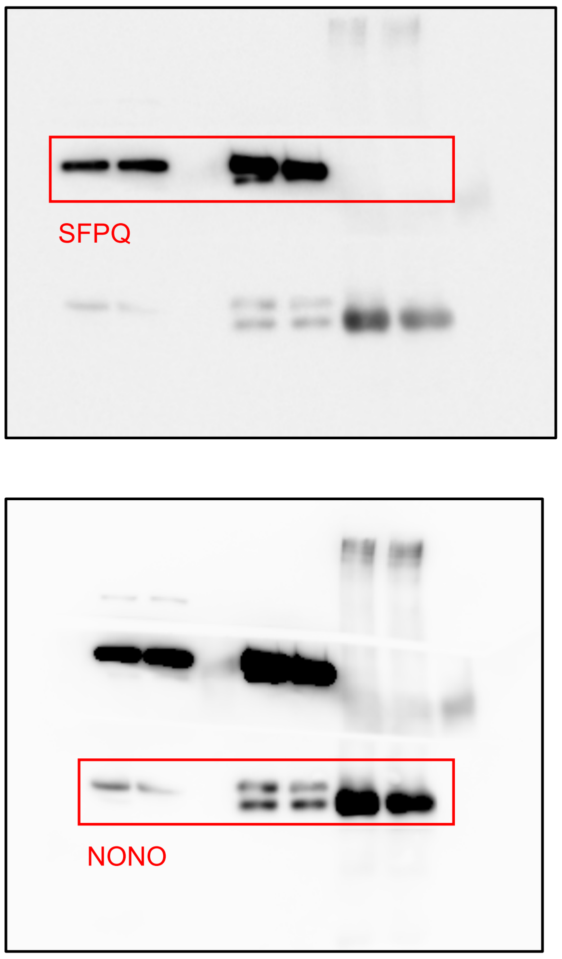


#### related to Fig. S1A

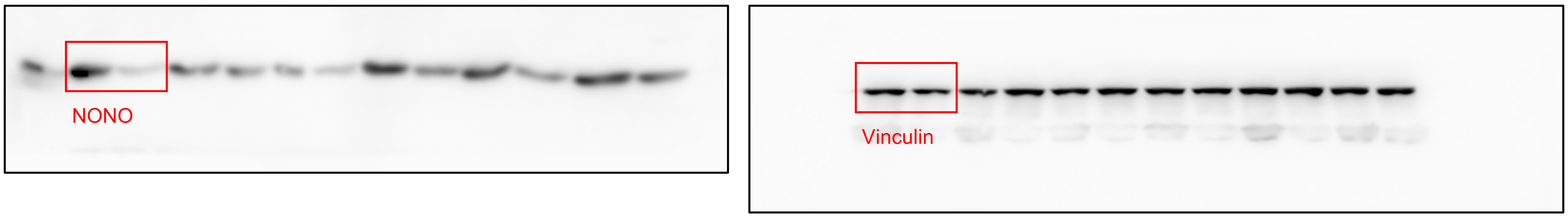


#### related to Fig. S1F

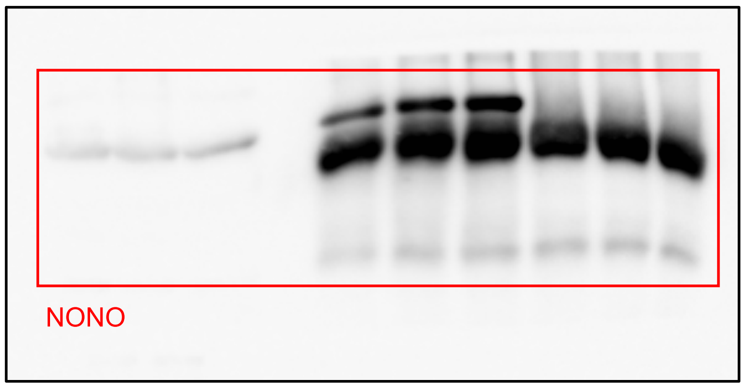


#### related to Fig. S3C

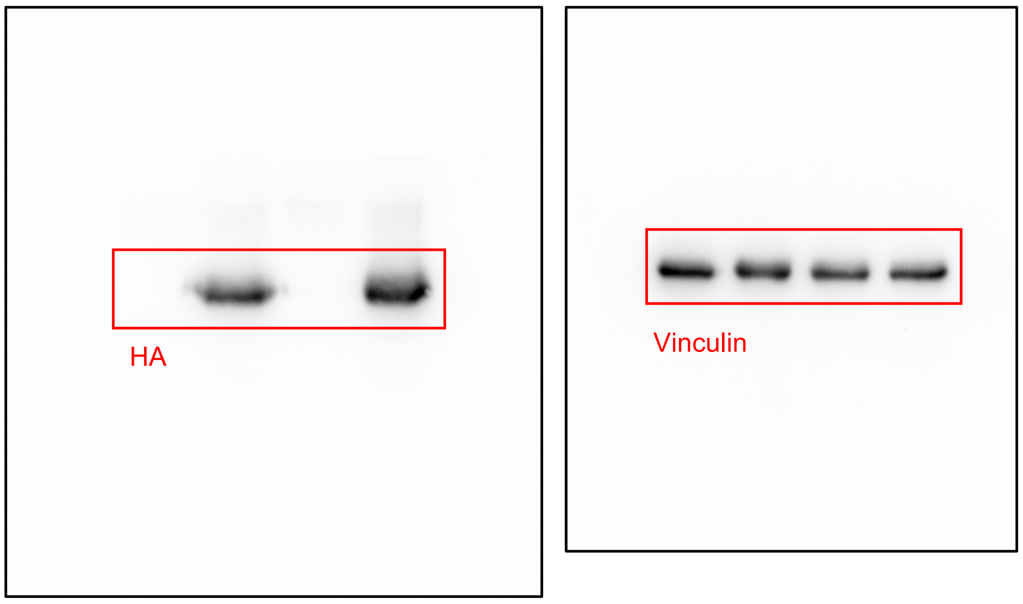


**Figure S4. Raw data.** Uncropped immunoblots are shown. Red box, cropped area depicted in figure panels.
